# Long-term impacts of tree monoculture plantations on biodiversity are mediated by soil acidification

**DOI:** 10.1101/2025.05.27.656426

**Authors:** Simone Balestra, Vanessa Manuzi, Erica Ceresa, Pietro Gatti, Reetta Aleksandra Pirttilahti, Thierry Adatte, Stephanie Grand, Gianalberto Losapio

## Abstract

Reforestation efforts have sought to counteract deforestation and to provide a nature-based solution against climate change. However, they often involve monoculture plantations of non-native species, which may have unintended ecological consequences. Yet, the long-term impacts of planting trees have been poorly estimated. Leveraging historical reforestation conducted in northern Italy during the 1920s by the fascist regime, we assessed the long-term impacts of red spruce (*Picea abies*) monoculture plantations on biodiversity of plants and soil fauna. We found that plant diversity in tree plantations was 50.3% lower than in native forests and 74.5% lower than in grasslands. Additionally, functional evenness was reduced by 30% in spruce plantations, suggesting lower ecological stability. In tree plantations, soil pH was significantly more acid and organic carbon content was 25% higher due to litter deposition and slower decomposition rates. Soil fauna diversity was marginally less affected, suggesting a faster recover over the last one-hundred years of arthropods as compared to plants. These findings highlight the need for monitoring reforestation interventions, suggesting strategies that incorporate diverse tree species rather than planting tree monocultures to support functionally and resilient ecosystems.

## 1 Introduction

Forests play a crucial role in supporting global biodiversity by providing habitats to 50–80% of all terrestrial organisms (Leuschner and Homeier, 2022). Furthermore, forests enhance ecosystem functions by contributing to regulating global climate via evapotranspiration and carbon cycling (Feng et al, 2024). Yet, deforestation and biodiversity loss continue to threaten the integrity and stability of forest systems (IPBES, 2019). Over the past few years, reforestation initiatives are increasingly implemented to mitigate forest losses and supposedly to mitigate the effects of climate change (Food and Agriculture Organization of the United Nations and United Nations Environment Programme, 2020). While reforesting and restoring degraded, deforested land is undoubtedly necessary for enhancing biodiversity recovery and mitigating climate change (Food and Agriculture Organization of the United Nations, 2020), unfortunately, at the moment, half of the area pledged for forest restoration is monoculture plantations characterized by non-native tree species (Lewis et al, 2019). Such plantations can have negative consequences for biodiversity, soil health, and ecological resilience (Messier et al, 2022) as timber production is maximized over biodiversity maintenance (Lewis et al, 2019). However, the long-term effects of reforestation on biodiversity remain poorly quantified.

Two key issues of are represented by (1) the poor species diversity and (2) the type and origin of trees used which involves non-native species. Such ‘reforestation’ should be named tree monoculture plantation. Native forests are rich and complex ecosystems able to provide a wide range of ecosystem services and nature’s contributions to people (Cavanagh and Benjaminsen, 2014). Yet, they are replaced by weak and poor monocultures that often display high mortality rates. On one hand, funding bodies tend to prefer material ecosystem services like timber provision over regulating services such as water retention. On the other hand, reforestation plans are often promoted to compensate for fossil fuel emissions by companies with large responsibilities for *CO*_2_ emissions such as airlines. Hence, few non-native and fast-growing trees like *Pinus* and *Eucalyptus* are vastly employed in plantations arrays worldwide (Uribe et al, 2021; Liang and Zong-Qiang, 2009). Recent studies (Feng et al, 2022; Grossman et al, 2018; Hua et al, 2022) reveal the benefits of multi-species plantations in terms of productivity, stability and biodiversity maintenance. Valuing plant diversity in reforestation programs may require a greater initial effort, but it would provide a much better outcome in terms of ecosystem health and services in the long term ((Liang et al, 2025). Monitoring the long-term consequences of reforestation for biodiversity is key for ensuring the wider sustainability of forest restoration actions.

Reforestation programs in Italy have been implemented at the beginning of the twentieth centuries as part of hydro-geological mitigation and recovery plans. The main drivers were associated with that socio-economic situation characterized by a fast demographic growth, the need of farmed land, energy resources and timber (Agnoletti et al, 2013). Furthermore, political drivers including the power of timber industry and the repression of farmers played a key role in plantations programs established by the fascist regime (Armiero et al, 2022). Criteria for choosing planting species were based on generalist and fast-growing species. In mountain areas of northern Italy, the species adopted in tree plantations was the red spruce (*Picea abies*). Spruce was also employed for its fast growth rate and its good timber quality, traits that promoted its widespread planting including in areas outside of his natural distribution range and habitats (Caudullo et al, 2016). The main consequence of these plantations is the creation of a new habitat dominated by a single non-native species. Red spruce is a non-deciduous tree that keeps a constant low light intensity in the understory throughout the year (Manuzi et al, 2024). Furthermore, red spruce has an impact on soil conditions by altering the humus layer and litter quality (Ranger and Nys, 1994; Augusto et al, 2002). This impact could influence soil chemistry and in turn could modify the community of plants and soil organisms. The soil-acidification effects of red spruce on soil could be exacerbated in the areas with alkaline, carbonate substrate, contributing to exclude native species adapted to more basic soil conditions. However, the impacts of red spruce plantations on biodiversity and soil conditions remain poorly monitored in the long-term. It is reasonable to hypothesize that spruce plantations impact biodiversity via two main pathways: directly, via removal of native forests in place of monoculture; and indirectly, via changing micro-environmental conditions such as soil acidity and nutrient availability.

The aim of this research is to assess how spruce monoculture plantations influenced biodiversity 100 years after planting. In addition, we tested the role of soil changes in the mediation of such impact. To address these questions and test these hypothesis, we investigated biodiversity changes of plant and arthropod communities across different habitats including native mixed forests and grasslands. We particularly focus on shifts in plant community composition in relation to ecological factors and changes in soil conditions.

## 2 Methods

### 2.1 Study area

We examined plant arthropod diversity as well as soil conditions in a set of permanent quadrants at two sites in the Italian Prealps during the growing season of the year 2023. The two chosen sites were (figure 1): Mount Bisbino (1300 m asl, Cernobbio, province of Como, Italy) and Alpe del Viceré (900 m asl, Albavilla, province of Como, Italy). Both sites rest on the Moltrasio limestone formation (Lower Jurassic) which is made by cherty limestones and marls. The bioclimate is temperate with warm and humid summers and fresh winters (Pesaresi *et al*., 2017); mean annual temperature is *c* 8.9 °C, mean summer temperature of *c* 18.3 °C, and annual precipitation is *c* 1350 mm/year. The natural vegetation is characterized by a deciduous forest dominated by *Fagus sylvatica* and *Acer pseudoplatanus* at Mount Bisbino site and by *Castanea sativa, Tilia cordata* and *Acer sp*. at the Alpe del Viceré site. Native forests and grasslands are sparse between reforested areas with red spruce. The studied grasslands have a mixed management that alternates between mowing and grazing.

**Fig. 1:**
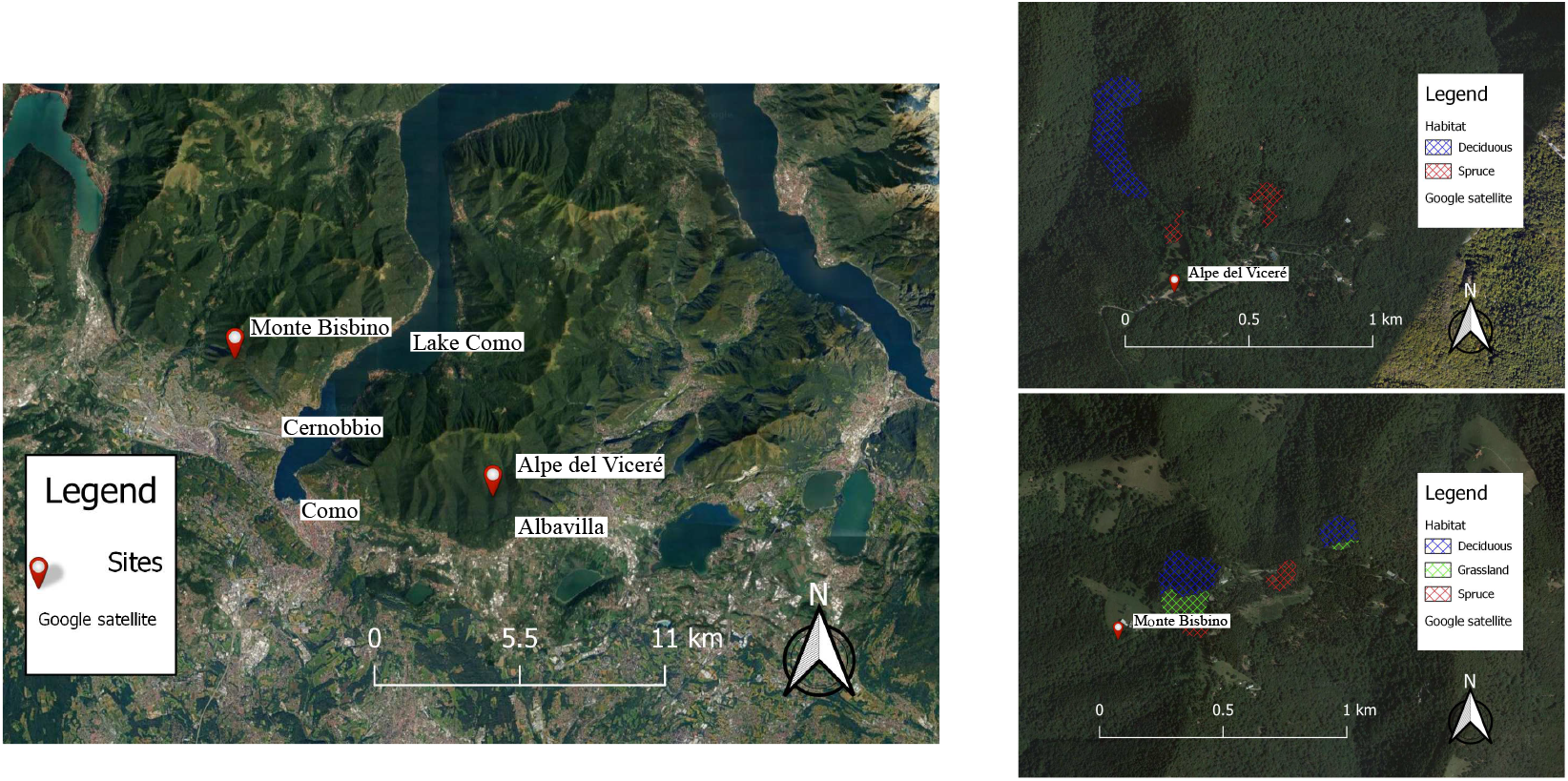
On the left the location of the sites studied. According to SOUISA classification (Marazzi, 2005), Alpe del Vicerè is part of the mountain chain “Triangolo Lariano” and Monte bisbino of “Tremezzo-Generoso-Gordona” chain, both in Como prealps (subsections of Lugano prealps). On the right the satellite images of the study sites. The colored patches represent the three habitats containing the fixed plots. In the legend “Spruce” stands for the spruce monoculture, “Deciduous” for the deciduous forest and “Grassland” is the mountain hay meadow.

Mount Bisbino plantations were installed during the 1930s (Bartolini, 2016) while Alpe del Viceré plantations were installed in 1930–1935 as part of the “Alpine Village” built by fascist regime, despite there were already present some spruce plantations in neighbouring areas (Borghese, 1992). In Alpe del Viceré site, it was not possible to sample the grasslands due to the local management of mowing and their occurrence at much higher elevation compared to the forests, conditions which would not provide the same bioclimate necessary for a robust comparison.

### 2.2 Field activity

We sampled plant and arthropod diversity belonging to the three habitats tested: spruce monoculture plantations, deciduous native forests and grasslands (Figure 1). We chose randomly five different plots for each vegetation of the two sites and marked a squared perimeter of 3 × 3 m, 9 *m*^2^ (n = 25). Every plot has been sampled monthly. We collected plant data taking account of the coverage provided by every species occurring in each plot. Only in spruce monoculture and deciduous forest, we sampled wood plants using a stratified method, which considers three levels of growth related to a specific stage of life history: seedling, sapling and adult. We sampled every plot monthly from March to July 2023 to better identify all species with different phenology. The species accumulation curve for sampling of plants is provided in Appendix A, showing the reliability of our sampling project.

We sampled the soil fauna community using a pitfall trap made with thermoplastic polyester (PLA) of 200 ml volume, filled with 50 ml of Propylene Glycol (C3H8O2). The traps were placed in the center of each plot, with the rim at the surface level, to attract mainly ground and litter arthropods. We made two cycles of arthropods samplings from April to June, letting the trap working for a week, for a total of 2 sampling rounds and 50 traps. Specimens were stored in an alcoholic solution (Et-OH 70%) prior to identification.

The species identification procedure followed different strategies. We identified plants according to “La Flora d’Italia” (Pignatti et al, 2017) and “Flora Alpina” (Aeschimann and Lauber, 2004). The arthropods have been observed and photographed at the stereo-microscope to detect specific the taxonomic traits. To identify arthropods, we created a project on the platform *iNaturalist* (https://www.inaturalist.org/projects/impact-of-spruce-plantation-on-arthropod-diversity) to take advantage of species identification provided by many experts around Italy and Europe. Overall, we identified 136 plant species and 253 arthropod taxa in total.

### 2.3 Soil analysis

Since we were interested in plant–soil interactions, we focused our soil analysis on the top two horizons: (1) the horizon 0, which is the organic one constituted by plant litter; (2) the horizon A, characterized by the organic mineral complexes. Two soil samples were taken from each plot, one per horizon, by pooling 100 g of samples from three random spots in the plot. Samples were air dried before the analysis. The samples have been sieved to remove the gravel particles bigger than 2 mm which are not considered as fine soil particles. We measured the pH of each sample following the protocol from (Pansu, 2006). We then analyze the content of the following elements:

- **Organic elements:** carbon, nitrogen, hydrogen.
- **Inorganic elements:** aluminium, magnesium, phosphorous, calcium, iron, potassium, sodium, manganese.

The organic elements Carbon and Nitrogen have been measured through an acid fumigation with concentrated HCl, which removes carbonates from calcareous soils (Harris et al, 2001). The other inorganic elements (Calcium, Magnesium, Potassium, Aluminium, Iron and Sodium) were measured following the multielement soil extraction known as “Mehlich 3 method” (Mehlich, 1984). We chose to consider for next step only the compounds that gave more meaningful results (C, N, K, Al, Fe and pH).

We kept those compounds that were expected to be more influenced by the spruce litter. The organic compounds carbon and nitrogen are a good indicator of the change of SOM turnover (Stevenson, 1994) and the metal free ions are expected to increase with soil acidification (Lundström et al, 2000). The outcomes for the compounds we did not evaluated are provided in the Appendix B.

### 2.4 Ecological indicators and indexes

Finally, to assess the long-term changes in ecological conditions, we adopted a Bioindication approach (Ivanova and Zolotova, 2023).

We assessed the Landolt’s Ecological Indicators (EIV) from (Landolt et al, 2010) and the related indexes and values for each plant species. These indicators ranges from 1 to 5 depending on the species’s ecology. We chose the following categories:

- **Light (L):** Refers to the plant’s tolerance and preference for light exposure. The scale typically ranges from species that thrive in deep shade (*L* = 1) to those that require full sunlight (*L* = 5).
- **Temperature (T):** Reflects the plant’s preference for specific temperature ranges. It ranges from cold-tolerant species occurring in alpine zone (*T* = 1) to those typical of warmer bioclimates (*T* = 5) than temperate ones.
- **Moisture (F):** Shows the plant’s preference for the moisture content of the soil. It ranges from very dry (*F* = 1) to waterlogged environments (*F* = 5).
- **Soil Reactions (R):** Indicates the soil reaction level that plants prefer. It ranges from highly acidic (*R* = 1) to ultra-alkaline (*R* = 5) soils.
- **Nutrients (N):** Indicates the plant’s tolerance or requirement for soil nutrients. It ranges from oligotrophic (*N* = 1) to eutrophic environments (*N* = 5).

We used the indicators to calculate the functional indexes through the R function “dbFD” from package FD (Laliberté et al, 2014). To this end, we built two matrices as input: one matrix with species in rows and EIV in columns, and a second matrix with plots in rows and plant species in columns. For functional analysis, we considered functional evenness (FEve). FEve measures how evenly the functional traits of a community are distributed, indicating how regularly the species fill the available functional space. (Mason et al, 2005; Laliberté and Legendre, 2010; Villéger et al, 2008). High FEve indicates an efficient and fair use of ecological niches, leading to a more stable ecosystem. On the contrary, a low FEve suggests that the species occupy only some areas of the functional space, with lacks in other traits. This could cause problems in resource use efficiency of the ecosystem.

### 2.5 Statistical analysis

We collected our data in different matrices and we have elaborated them with RStudio 2023.12.1 (Posit Team, 2023), using the packages mixOmics (F et al, 2017), lavaan (Rosseel, 2012), parameters (Lüdecke et al, 2020), effectsize (Ben-Shachar et al, 2020), rfPermute (Archer, 2023) and vegan (Oksanen et al, 2022). First, we performed regression analysis to assess how biodiversity and soil change with spruce monoculture as compared to native forests and grasslands. In particular, for plant and arthropod diversity, we tested the response of species richness (two separate models for plants and arthropods) to spruce monoculture (reference level, with native forests and grasslands as conditional levels) using a generalized linear mixed model (GLMM) with a Poisson distribution and site as a random effect. For EIV and FEve, we used linear mixed models (LMM) for each soil parameter (Light, Temperature, Moisture, Soil Reactions and Nutrients) including spruce monoculture (reference level, with native forests and grasslands as conditional levels) as fixed effect and site as random effects. For soil conditions, we used LMM for each soil parameter (carbon, nitrogen, aluminium, iron, potassium and pH) including spruce monoculture (reference level, with native forests and grasslands as conditional levels) and soil horizon (categorical with horizons 0 and A) as fixed effects, including their statistical interactions too, and site as random effects.

Second, we used multivariate analysis, specifically Partial Least Square Discriminant Analysis (PLSDA)and a path analysis. We did PLSDA to study the plant communities of the sites, with a focus on the differences between the monoculture and the natural habitats. PLSDA was performed on a binary matrix with the presence/absence of species distribution as predictors. In this way we used the species to discriminate among the habitats, identifying also which ones contribute more to discriminate one habitat from the others. Path analysis was developed starting from a stepwise model with selection based on AIC criterion and the backward elimination, as done by (). We selected two equations with plant diversity and arthropod diversity as dependent variables, since we wanted to see how our variables could affect biodiversity. We built each equation by putting all the variables that was potential predictors for us. Both for plants and arthropods we used the soil parameters (carbon, nitrogen, aluminium, iron, potassium and pH), the ecological indicators (Light, Temperature, Moisture, Soil Reactions and Nutrients), a binary variable of habitat naturalness (1 stands for “monoculture” and 0 for “natural habitat”) and a binary variable of habitat structure (1 for “herbaceous” and 0 for “forest” habitat). For arthropod diversity, we considered also plant diversity as predictor, since plants are a source of food for many arthropod taxa. The stepwise selection have removed the redundant variables, in order to obtain the lowest AIC value, which indicates the best fit for our model (Yamashita et al, 2007).

## 3 Results

### 3.1 Biodiversity

Plant diversity was found to be strongly dependent on vegetation (*p* − *value <* 0.001), with a clear trend of species increasing from spruce monoculture to grassland (Figure 2 a). In spruce monoculture plant diversity is 50.3 % lower than deciduous forest and 74.5 % lower than grassland. The median of species per plot was 7 for monoculture, 18.5 for deciduous forest and 37 for grassland. Arthropod diversity model turned out as non-significant (*p* − *value* = 0.24) though the graphic (figure 2 b) shows a trend of increase from spruce monoculture to grassland. The median of arthropod taxa per plot was 25 in the monoculture, 28.5 in deciduous forest and 37 in grassland. The IQR of deciduous forest has shown a very high data dispersion.

**Fig. 2:**
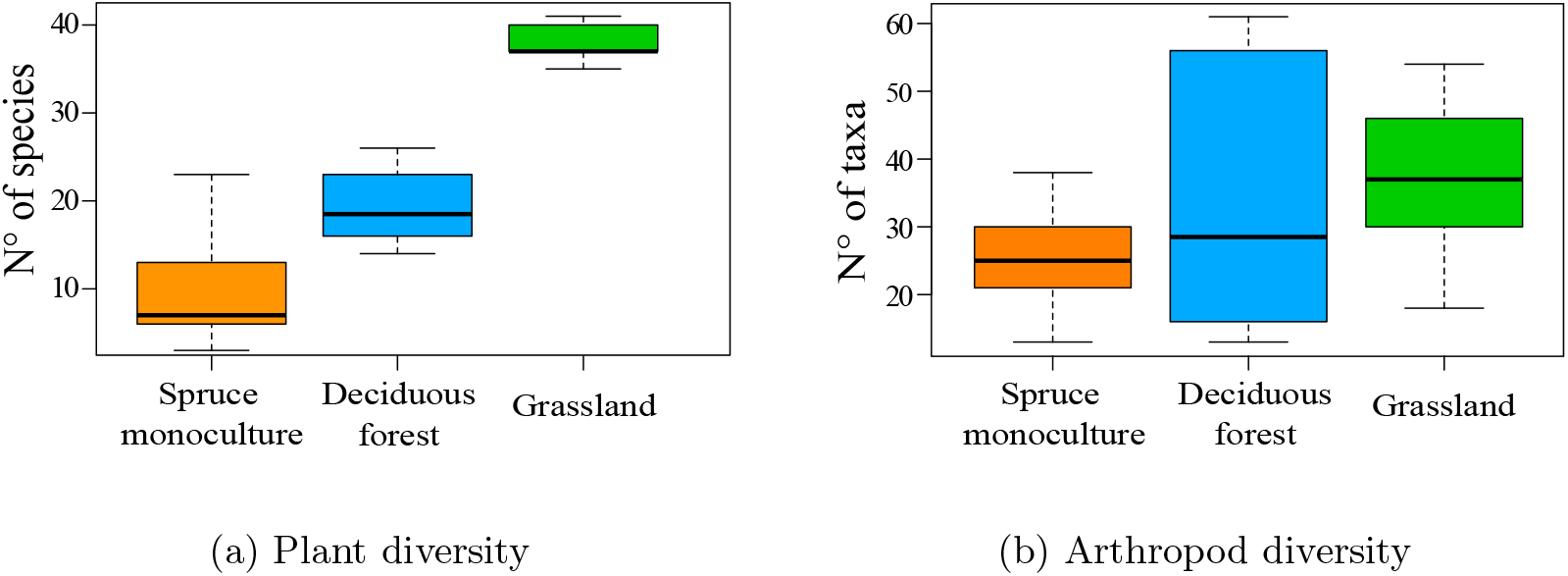
Box plots representing how plants and arthropod diversity change among habitats. On the x-axis are reported the three habitats, on the y-axis the diversity is represented as number of species. The colorful rectangle represent the IQR (Interquartile Range), that is the central 50 % of data in their distribution. The bold black line represent the median that divides the box in first quartile (lower half) and third quartile (hupper half). At the ends of the box there are the whiskers, which indicate the data dispersion outside the IQR without include the outliers.

### 3.2 Partial least square discriminant analysis

We produced two graphs: first, a Scatter plot that shows how the plots are distributed in the space of components; second, a dendrogram that highlights the discriminant species among the habitats.

The scatter plot (figure 3a) displays the plots in the components space. The first and second components show significant differences in species distribution among spruce monoculture and the natural habitats. The statistical tests driven turned out to be significant for both first (*p* − *value <* 0.001) and second (*p* − *value* = 0.005) components. The first component explained 23 % of variance, distinguishing the three habitats from each other. In particular, it shows a strong differentiation among the grassland (grey ellipse) and the forests (blue and orange ellipses). The second axis explained 7 % of variance, differentiating deciduous from and spruce monoculture.

**Fig. 3:**
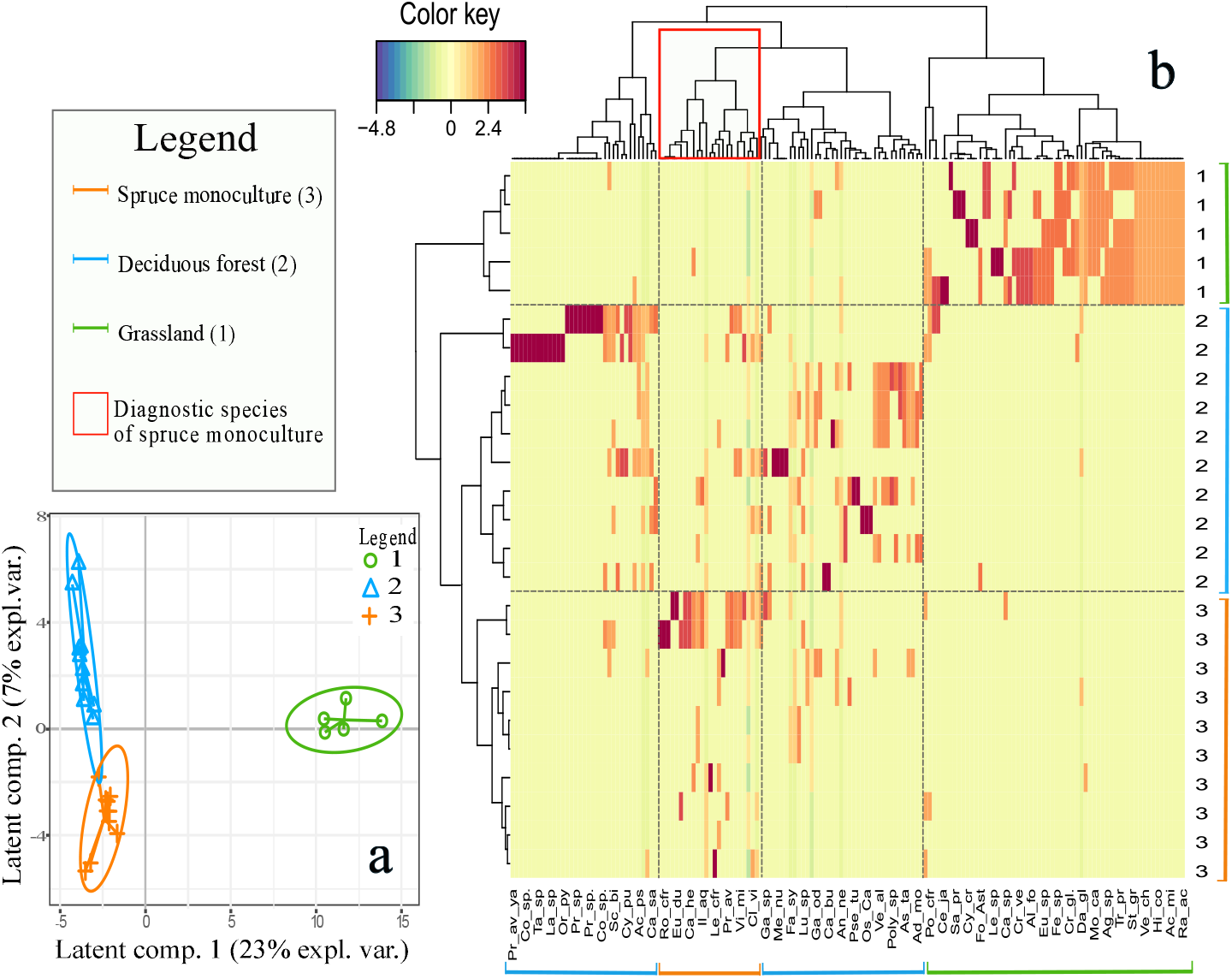
Graphs resulting from the PLS-DA analysis on the plant community. At the bottom left: Scatter plot representing the distribution of the plots in the space of the components identified by the model; along the two axes is expressed the percentage of variance explained by the model. In the remaining part of the figure: Dendrogram and legends referring to it, representing the differentiation of the plants sampled. On the bottom of the graphic are listed the species identified by the model to distinguish habitats from each other. On the right side are listed the plots classes. The color key indicates the grade of differentiation among habitats and shows an increase of intensity from blue to red. In the dendrogram on the top side is highlighted the cluster of species that best distinguish the monoculture from the other habitats.

The graphic with dendrograms (Figure 3b) deepens the result found in the previous one. The mutual differentiation of habitats in the first component is represented in the left dendrogram, where the upper branch well distinguish the grassland from the two forests. Results of PLS-DA provide a set of plant species that best differentiate habitats (Figure 3 Table 1). The upper dendrogram shows how the species are clustered depending on their presence/absence in the habitat. This dendrogram again distinguishes very well the pool of species related to grassland from the one of the forests. It also provides a more precise picture of how the plant community of spruce monoculture is related to the one of deciduous forest. the plant species of spruce monoculture form a cluster (red rectangle in Figure 3b) included in the same hierarchical level of the clusters related to deciduous forest; and does not belong to a separate cluster as one would expect in case of habitat differentiation.

**Table 1:** species automatically identified by the PLS-DA model as distinguishing factors of the plant communities.

| Spruce monoculture | Deciduous forest | Grassland |
| --- | --- | --- |
| *Primula vulgaris* | *Corylus avellana* | *Arrhenatherum elatius* |
| *Geranium robertianum* | *Acer pseudoplatanus* | *Ranunculus acris* |
| *Cardamine bulbifera* | *Veratrum album* | *Thymus sp.* |
| *Daphne laureola* | *Castanea sativa* | *Achillea millefolium* |
| *Prunus avium* | *Ranunculus platanifolius* | *Cruciata glabra* |
| *Rosa sp.* | *Lilium bulbiferum* | *Lathyrus pratensis* |
| *Geranium nodosum* | *Narcissus poeticus* | *Hippocrepis comosa* |
| *Brachypodium sylvaticum* | *Lonicera xylosteum* | *Betonica officinalis* |
| *Urtica dioica* | *Adoxa moschatellina* | *Galium lucidum* |
| *Larix decidua* | *Sorbus aria* | *Veronica chamaedrys* |
| *Hieracium sylvaticum* | *Geranium nodosum* | *Potentilla erecta* |
| *Helleborus viridis* | *Asperula taurina* | *Hieracium pilosella* |
| *Ilex aquifolium* | *Senecio nemorensis* |  |

### 3.3 Ecological indicator values (EIVs)

#### 3.3.1 EIVs estimation

We found that EIVs of soil pH show no significant differences between spruce monoculture and other habitats (*p* − *value* = 0.86, Figure 4a). The values for spruce monoculture have a high variance, suggesting the presence of a wide range of plant species adapted to grow in different soil pH conditions. The EIVs of temperature indicate that deciduous forest and grassland provide habitats for communities with higher values than the spruce monoculture (*p* − *value* = 0.001; Figure 4b). Species related to spruce appear to be well adapted to colder climatic conditions. The EIVs of humidity indicate that spruce monoculture provides a habitat with no difference than deciduous forest. The two forests have both higher values than grassland (*p* − *value* = 0.004; Figure 4c). Species related to spruce monoculture and deciduous forest seem to be adapted to higher values of humidity. The EIVs of root depth indicate that species of spruce monoculture have deeper roots than deciduous forest and grassland (*p* − *value <* 0.001; Figure 4d). The decreasing trend suggests that the species of the three habitats are mutually different for this ecological indicator and the lowest value is in plant community of grassland. The EIVs of luminosity indicate that spruce monoculture provides habitat for communities with lower values than the deciduous forest, which in turn has lower values than grassland (*p* − *value <* 0.001; Figure 4e).

**Fig. 4:**
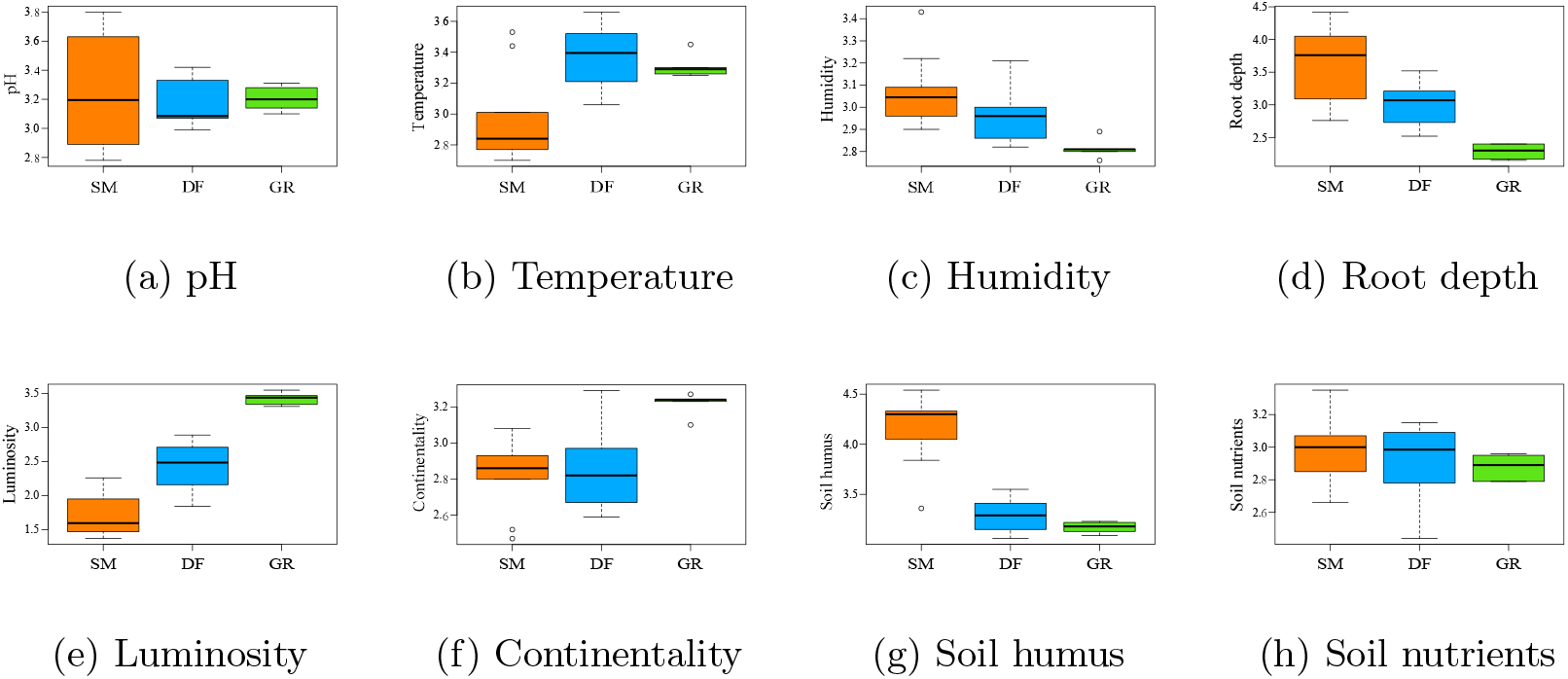
Box plots representing how the ecological indicators from (Landolt et al, 2010) assigned to the plants species sampled changes among different habitats. (a) ranges following the classic scale (acid to basic). (b) ranges from low to high. (c) ranges from dry to humid. (d) ranges from low to high soil loose. (e) ranges from shadow to light environment. (f) ranges from oceanic to continental climate. (g) is referred to humus content. (h) is referred to nitrogen content.

The species of spruce monoculture are better adapted to lower level of luminosity than species of deciduous forest and grassland. The EIVs of continentality indicate that the spruce monoculture (and deciduous forest as well) have a pool of species with a lower value of continentality compared to grassland (*p* − *value* = 0.002; Figure 4f). The plant community of grassland have higher values, meaning that its species are well adapted to climates with low precipitation and higher thermal amplitude. The EIVs of soil humus suggests that spruce monoculture offers a habitat with much higher amount of soil humus than deciduous forest and grassland (*p* − *value <* 0.001; Figure 4g). The species of monoculture are therefore expected to be well adapted to soil with slow turnover and high accumulation of organic matter. The EIVs of soil nutrients do not show differences among the three habitats (*p* − *value* = 0.41; Figure 4h). The adaptation to a certain nitrogen content in soil for the species of spruce monoculture seem not to differ from deciduous forest and grassland. The additional information on model statistics are resumed in Figure 4, Table 2.

**Table 2:** Table of statistical parameters from Figure 4. The column represent: model p-value, p-value of habitat comparison (DF = Deciduous Forest, GR = Grassland and SC = Spruce Monoculture) and R^2^. The rows are the Ecological Indicators, listed in the same order of Figure 4.

| EI | Model p | BM vs FC (p) | PR vs FC (p) | R<sup>2</sup> |
| --- | --- | --- | --- | --- |
| pH | 0.861 | 0.590 | 0.798 | 0.0135 |
| Temperature | 0.00125 | 0.00047 | 0.01011 | 0.4554 |
| Humidity | 0.00349 | 0.05768 | 0.00094 | 0.4021 |
| Root Depth | <0.00001 | 0.00214 | <0.00001 | 0.6274 |
| Luminosity | <0.00001 | 0.00002 | <0.00001 | 0.8354 |
| Continentality | 0.00185 | 0.84024 | 0.00089 | 0.4356 |
| Soil Humus | <0.00001 | <0.00001 | <0.00001 | 0.7927 |
| Soil Nutrients | 0.4084 | 0.2950 | 0.2390 | 0.0782 |

#### 3.3.2 Functional evenness

The functional evenness returned meaningful results, useful to define every habitat type as the overall model was significant (*p* − *value* = 0.014; figure 5). In particular, functional evenness of spruce monoculture was 30% lower than both deciduous forest (*p* − *value* = 0.009) and grassland (*p* − *value* = 0.02). Deciduous forest and grassland are also significantly different, despite they have a small differences (*p* − *value* = 0.019 and *estimate* = 0.092).

**Fig. 5:**
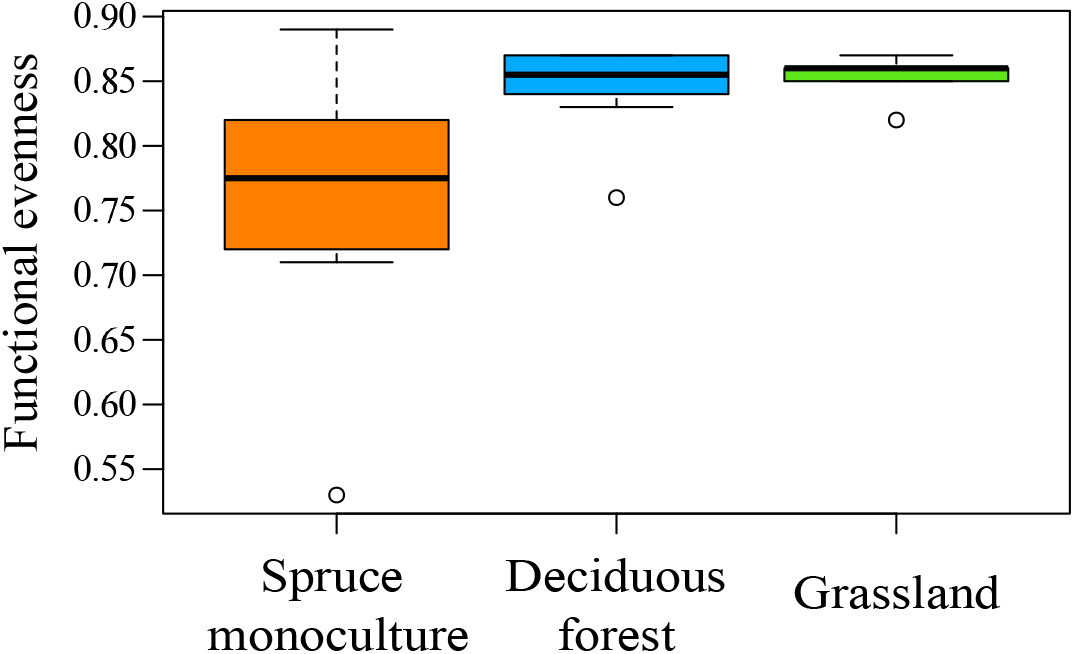
Box plots representing the levels of the ecological indicator functional evenness in the three different habitats.

### 3.4 Soil conditions

The results (figure 6) show differences in soil properties between both habitats and horizon level. pH (6a) has shown expected results, being significant for vegetation (*p* − *value* = 0.048) with more acid values in spruces monoculture. No significant differences among horizons have been detected. Both vegetation (*p* − *value <* 0.001) and horizon (*p* − *value <* 0.001) have an effect on Carbon (6b), with higher values in the “0” than the “A” horizon and a trend of decreasing from the spruce monoculture to the grassland. This result is in accordance with the “Soil humus” parameter in previous section (par. 3.3.1, Figure g); and indicates a higher accumulation of organic matter in the spruce forest understory. Nitrogen (6c) changes only among horizons (*p* − *value <* 0.001), where shows higher values in the “0”.

**Fig. 6:**
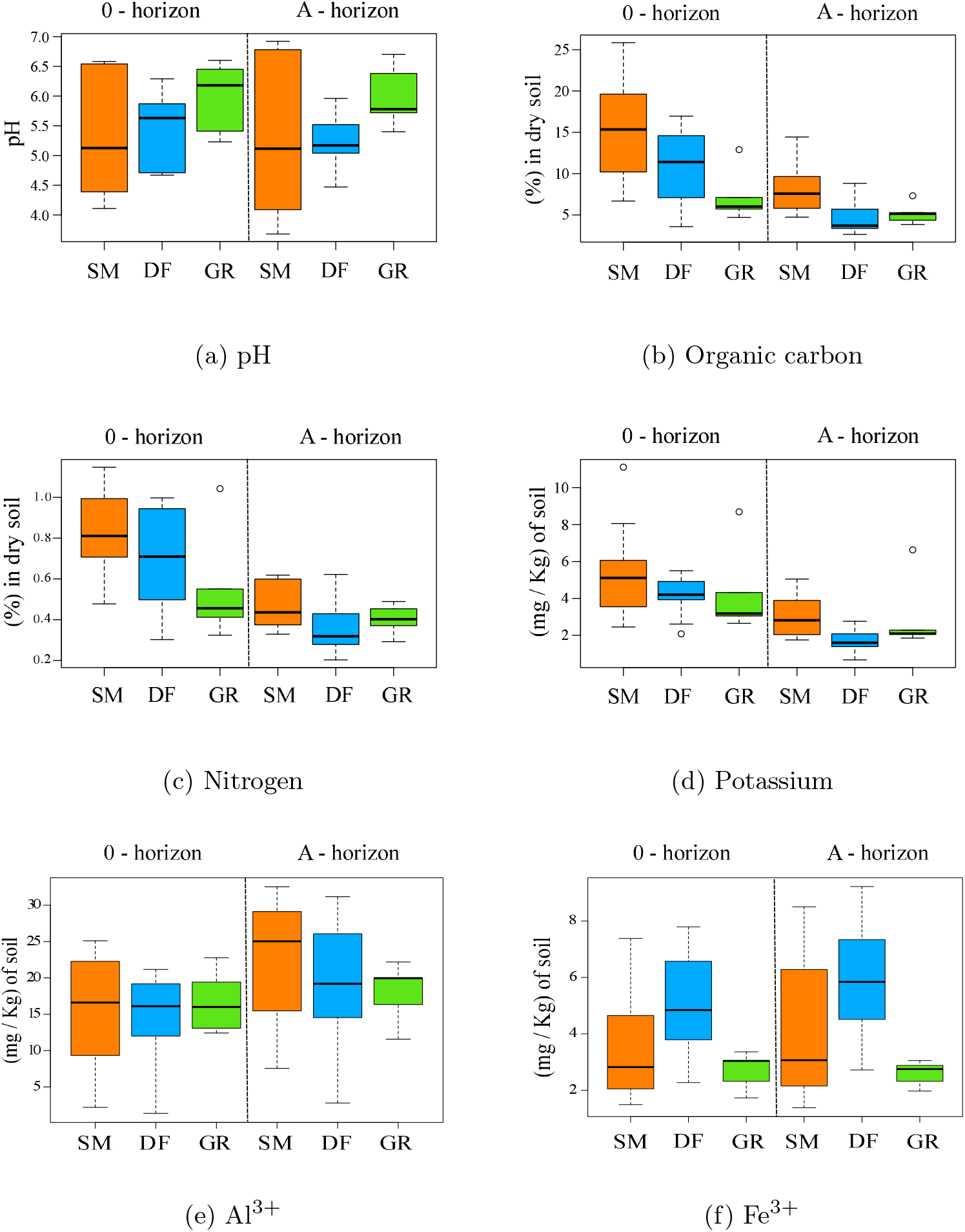
Box plots representing the differences of soil parameters for the three habitats, divided by horizon. On the x-axis are indicated the habitats (SM = spruce monoculture, DF = deciduous forest and GR = grassland). On the top of graphic are indicated the soil horizons (“0” horizon = litter, “A” horizon = organic mineral complexes). The y-axis shows the scales of each parameters taken into account.

Although the relationship was not significant (*p*−*value* = 0.059), the graphic shows slightly higher nitrogen values in the monocultured spruce soil. Vegetation changes Potassium (6d) values (*p* − *value* = 0.01) with the same trend of (6b) and horizon “0” has higher levels (*p* − *value <* 0.001). The free *Al*^3+^ (6e) was higher in “A” horizon (*p* − *value* = 0.01). Despite there is a trend of decrease in aluminum from spruce monoculture to grassland the differences with natural habitats were not significant. The free *Fe*^3+^ (6f) had no differences among both the two horizons and the three habitats.

### 3.5 Path analysis

The results of the stepwise selection with backward elimination are reported in the following functions (1) and (2).

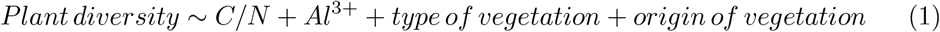

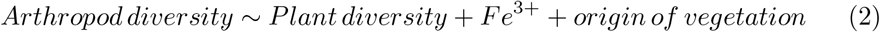

- **C/N** is the ratio of the SOC Carbon and Nitrogen. We combined these two parameters in a ratio according to their ecological meaning as indicators of nutrients recycling rate in soil (Gundersen et al, 1998; Ollinger et al, 2002).
- ***Al***^**3+**^ and ***Fe***^**3+**^ are the inorganic elements Aluminium and Iron.
- **“*type of vegetation*”** is a binary variable where 1 stands for “herbaceous” and 0 for “forest” habitat.
- **“*origin of habitat* “is** a binary variable where 1 stands for “monoculture” and 0 for “natural habitat”.

The good fit of the model have been reported in the following table (Figure 7 Table 3). We evaluated it considering the parameters suggested by Kline (2023) and Lomax (2004). The RMSEA was slighlty over the cut-off value, that indicates an unremarkable goodness of fit. However, this indicator can be misleading in model with few degrees of freedom (Kenny et al, 2015), as our. In addition, all the others indicator returned a good fit.

**Table 3:** Comparison of obtained values with acceptable cut-off criteria for goodness-of-fit indices. The first row indicates the threshold.

| Parameter | $p(\chi^2)$ | GFI | AGFI | NFI | NNFI | CFI | RMSEA | SRMR | RFI |
| --- | --- | --- | --- | --- | --- | --- | --- | --- | --- |
| Threshold | $> 0.05$ | $\geq 0.90$ | $\geq 0.90$ | $\geq 0.90$ | $\geq 0.95$ | $\geq 0.95$ | $< 0.08$ | $\leq 0.08$ | $\approx 1$ |
| Value | 0.307 | 0.996 | 0.974 | 0.933 | 0.963 | 0.987 | 0.090 | 0.040 | 0.817 |

**Fig. 7:**
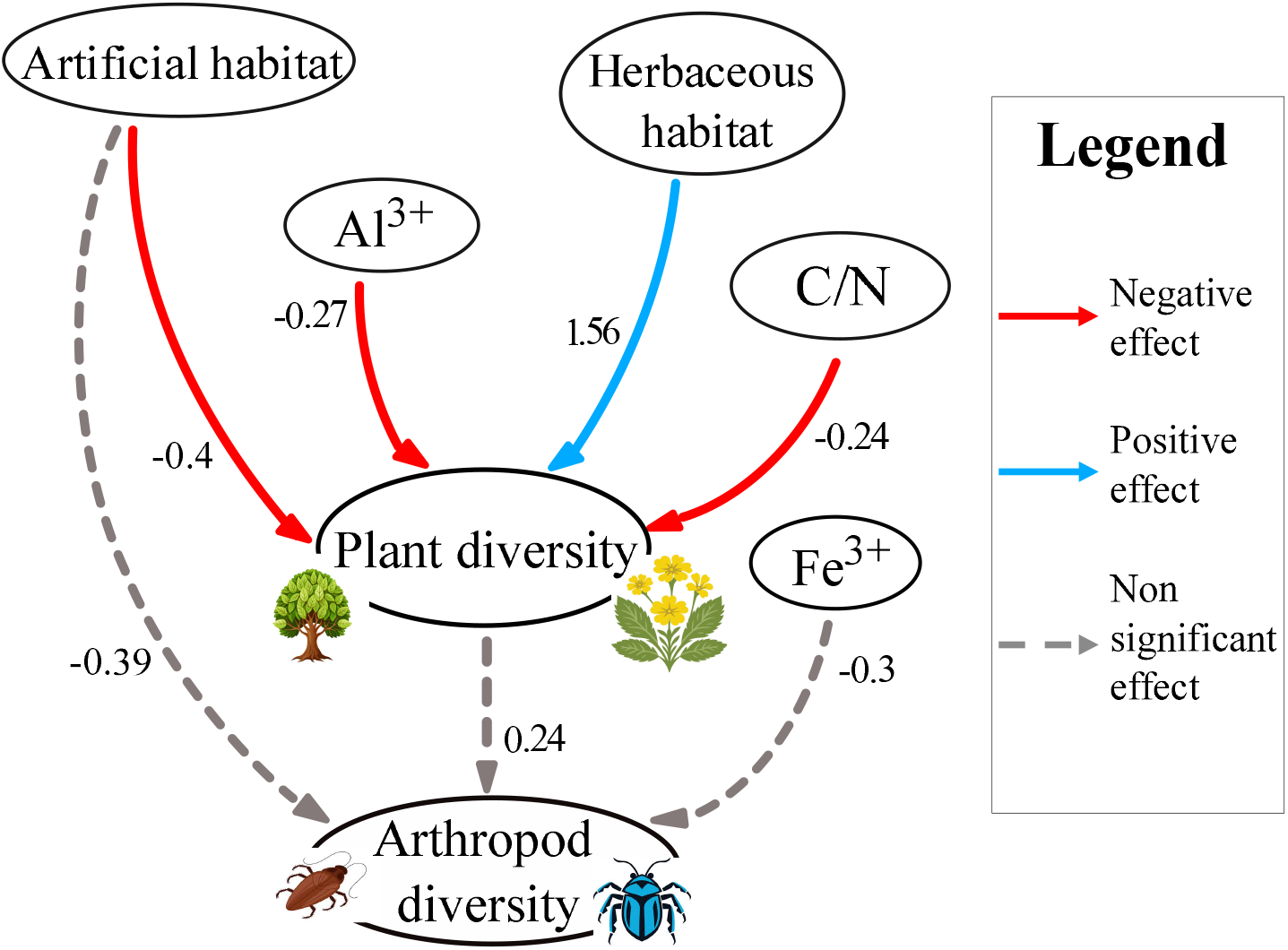
Path diagram resulting from the path analysis. The arrows represent the direct effect of a variable, labeled with the estimated coefficient value. The nodes contain the variables.

The path diagram (Figure 7) returned the graphical frame of the predictor’s effect on the two dependent variables. Plant diversity has shown evidence of undergoing significant effects from soil parameters and habitat properties. Higher levels of Al^3+^ and C/N have a negative effect on plant diversity. The herbaceous structure implies an higher plant diversity, with a strong coefficient. The anthropic origin of the habitat have a negative impact on it. Arthropod diversity does not appear to be influenced by the variables filtered by the model. Plant diversity does not have a significant effect on arthropod diversity as well, despite we considered it as a potential strong predictor.

This result is consistent with what was obtained in the analysis of biodiversity (paragraph 3.1), where no differences in arthropod diversity among spruce monoculture and natural habitats occurred.

## 4 Discussion

To address our hypotheses, we evaluated how spruce monoculture differs from natural habitats in terms of plant and arthropod biodiversity, functional indices of plant communities, and soil conditions. We found that spruce monoculture alters plant community composition, reduces plant diversity and functional evenness, and modifies soil conditions, particularly by reducing light availability and increasing pH and organic matter. Arthropod diversity was similar to native forests and grassland habitats.

### 4.1 Plant and arthropod diversity

Our comparative approach helped clarify how plant and arthropod communities respond to spruce monoculture plantations in the long-term. As expected, grasslands hosted the highest plant diversity. Less predictable, however, was the observed difference between the two forest types. This may be explained by the absence of early-flowering species and the absence of other tree species in spruce monoculture. These species are typical of deciduous forests, where they bloom before canopy closure in spring. The evergreen nature of spruce prevents such phenological adaptation, likely reducing species richness. (Hérault et al, 2005) The lack of tree species recruitment might be due to allelopatic effects associated with soil acidification and litter deposition (Aarrestad et al, 2014). While plant diversity was clearly habitat-dependent, arthropod diversity showed higher variation within habitats than between habitats. This result may be due to multiple factors. First, arthropod are mobile organisms that, in the absence of dispersal barriers, could easily move among the examined adjacent habitats (Anderson et al, 2024). Second, arthropod diversity is not exclusively structured by habitat but also by trophic relationships, including plant diversity (Losapio et al, 2016; Ebeling et al, 2018) A more diverse plant community typically supports a wider range of arthropods by providing more niches and food sources (Scherber et al, 2010). However, arthropods are highly adaptable and can exploit a broad range of trophic resources (Eisenhauer et al, 2019). Moreover, our sampling focused primarily on soil-dwelling arthropods, whose diversity might be more closely tied to biomass productivity and soil structure than to plant diversity. Yet, the positive trend observed in the arthropod model suggests that increased sampling intensity could yield clearer patterns.

### 4.2 Partial least square discriminant analysis

The PLS-DA confirmed distinct plant community compositions across habitats, validating the presence of three ecologically differentiated systems. Grasslands formed a distinct group, consistent with their unique structure and dominance of herbaceous species intolerant to shade. The closer proximity of the deciduous forest and spruce monoculture ellipses reflects their shared structural features and overlapping species pools. Further analysis revealed that the species diagnostic of spruce monoculture are nested within a broader group typical of deciduous forests. This suggests that spruce monoculture represents a taxonomically impoverished subset of the native forest community, consistent with our hypothesis that monoculture reduces local plant biodiversity. Importantly, the species distinguishing native, deciduous forest from spruce monoculture (e.g., *Oxalis acetosella, Corylus avellana*) are not exclusive to deciduous systems and may also occur in coniferous forests. Thus, the absence of distinct diagnostic species in spruce monoculture underscores its lower ecological integrity and biodiversity value.

### 4.3 Ecological indicator values (EIVs)

Landolt EIVs provided further insights into habitat characteristics. The pH indicator showed that soil pH alone does not define plant communities, although greater variance in spruce monoculture suggests it supports a mixed assemblage of species. Higher values for root depth may reflect the specific rooting strategies associated with spruce, maybe influenced by his presence in a warmer climate (Mauer et al, 2009). Similarly, the humidity index highlighted the influence of forest canopy. Both forest types exhibiting lower light availability than grassland, that enhanced protection from water loss due to evapotranspiration and subsequent desiccation (Liu et al, 2022). Luminosity was highest in grassland and lowest in spruce monoculture due to the evergreen canopy, which limits light penetration year-round (Stenberg et al, 1999). This likely restricts early-flowering geophytes, which are prevalent in deciduous forests. The absence of these light-dependent species in spruce monoculture reduces temporal niche availability and contributes to lower diversity (Manuzi et al, 2024). The continentality index was highest in grassland, consistent with species typical of continental climates. Spruce monoculture exhibited lower temperature values, possibly reflecting the cooler microclimate associated with coniferous canopy cover and higher-altitude species (Brna et al, 2024). Higher soil humus values in spruce monoculture align with reduced organic matter turnover. Nutrient indicators (e.g., nitrogen) did not distinguish habitats, suggesting similar levels of nitrogen utilization across plant communities.

Functional evenness differed significantly among habitats. Following Mason et al (2005) and Villéger et al (2008), functional evenness reflects the extent to which a community uses the available niche space. Lower values indicate that parts of the functional space are under-utilized, potentially reducing ecosystem productivity and resilience. Spruce monoculture’s low functional evenness suggests inefficiencies in resource use and greater vulnerability to disturbance. This may be due to spruce’s role in altering soil pH or to the exclusion of certain functional groups, such as early-flowering geophytes.

### 4.4 Soil conditions

Soil analysis confirmed that spruce monoculture alters soil composition, particularly in terms of soil organic carbon, nitrogen, and potassium, indicating slower organic matter turnover. The higher retention of organic matter in forests (compared to grasslands) likely reflects both accumulation and reduced decomposition rates (Liao et al, 2006). Although we did not observe reduced arthropod diversity in spruce monoculture, the slower turnover may be related to microbial communities, which were not assessed but may be less diverse in monoculture due to reduced plant inputs and microhabitat heterogeneity. Soil pH in spruce monoculture was lower, likely due to acidic needle litter that promotes podzolization and mineral leaching (Lundström et al, 2000). In contrast, grassland pH reflects the buffering effect of carbonatic parent material, which may counteract acidification processes, especially on sloped terrain (Borvka et al, 2007). Both the free Al^3+^ and Fe^3+^ was not higher in “0” horizon of spruce monoculture, as we could expect as consequence of soil acidification drived by spruce litter. However, the trend of Al^3+^ suggests a potential significant variation between spruce monoculture and natural habitats, might be achieved with a greater number of samples.

### 4.5 Path analysis

The effect of Al^3+^ and C/N on plant diversity have a direct relation with the impact of spruce on soil development. Both organic carbon and nitrogen two soil parameters have been found higher in the monoculture compared to natural habitats (paragraph 3.4). While for Al^3+^ the higher amount in spruce monoculture was not statistically significant. High C/N ratio could mean a low turnover of organic matter, creating a condition of low nutrient availability. High Al^3+^ is associated to acid soils with strong chemical erosion of parent material (Lundström et al, 2000) and may act as an ecological limitation for some plant species. Arthropod diversity was not influenced by any predictor. This result, in accordance with what we observed in Figure 2b, suggest that the arthropods are not negatively influenced by the spruce. This could be due to their high adaptability and mobility, as we discussed in paragraph 4.1. As we hypothesized, the presence of an artificial habitat have a negative effect on plant diversity of the study area. This confirms that the presence of a species as red spruce that has a strong impact on the landscape due to its dominance, outside his optimal range, can be detrimental for biodiversity.

## 5 Conclusion

This study case demonstrates the long-lasting impacts of planting trees on biodiversity. Specifically, it highlights that spruce monoculture negatively impacts plant biodiversity and functional evenness, primarily through changes in soil conditions (pH, organic matter, and humus accumulation). Arthropod diversity appeared less sensitive to habitat differences, possibly due to higher mobility and quicker recovery. Taken together, we revealed that spruce monoculture is essentially a degraded subset of the deciduous forest community, contributing little to overall biodiversity. Soil parameters further distinguish spruce monoculture, highlighting the effects of needle litter on soil development, though some effects may be mitigated or exacerbated by underlying geology. Landolt EIVs effectively captured ecological differences among habitats, with spruce monoculture favoring species adapted to high humus content, low light, and low temperature. Functional evenness results suggest that spruce monoculture is less stable and efficient, which may compromise long-term ecosystem functioning.

Our study illustrates the ecological consequences of reforestation strategies that prioritize canopy cover over biodiversity and soil conditions. Spruce alters key environmental conditions and limits community functional diversity. Even native species, when planted outside their optimal ecological range, can significantly affect local ecosystems. This insight has broader implications: if native spruce can degrade biodiversity, the risks posed by fast-growing, non-native species in global reforestation efforts may be even greater. Understanding the long-term impacts of past monocultures can inform current and future reforestation programs to ensure that ecological restoration promotes both biodiversity and ecosystem functioning while mitigating the the impacts of climate change.

## Author contributions

Conceptualization: S.B., G.L.; Data collection: S.B., V.M., E.C., P.G., R.A.P.; Labwork: S.B., V.M., E.C., P.G., T.A., S.G., Data analysis: S.B., G.L.; Writing—original draft, S.B.; Writing—review & editing: S.B., G.L.; Funding acquisition: G.L.

## Data availability

Data will be made publicly accessible on Zenodo. The R code to analyse data and to reproduce the results and figures reported in this manuscript will be publicly accessible on GitHub.

## Declarations

### Competing interests

The authors declare no competing interests.

## 6 Appendix

### 6.1 A - Species accumulation curves

The species accumulation curves of the three habitats (Figure 8) have confirmed an adequate sampling effort. The amount of species sampled in the curves of spruce monoculture and deciduous forest tend to reach a plateau prior to the total amount of plots on the x-axis. The curve of grassland instead, seem to reach the beginning of the plateau at the last plots. This lower (but still acceptable) sampling efficiency in grassland is due to the lower number of plots we made in grassland. As we already mentioned (par. 2.2), this shortage was due to local management.

**Fig. 8:**
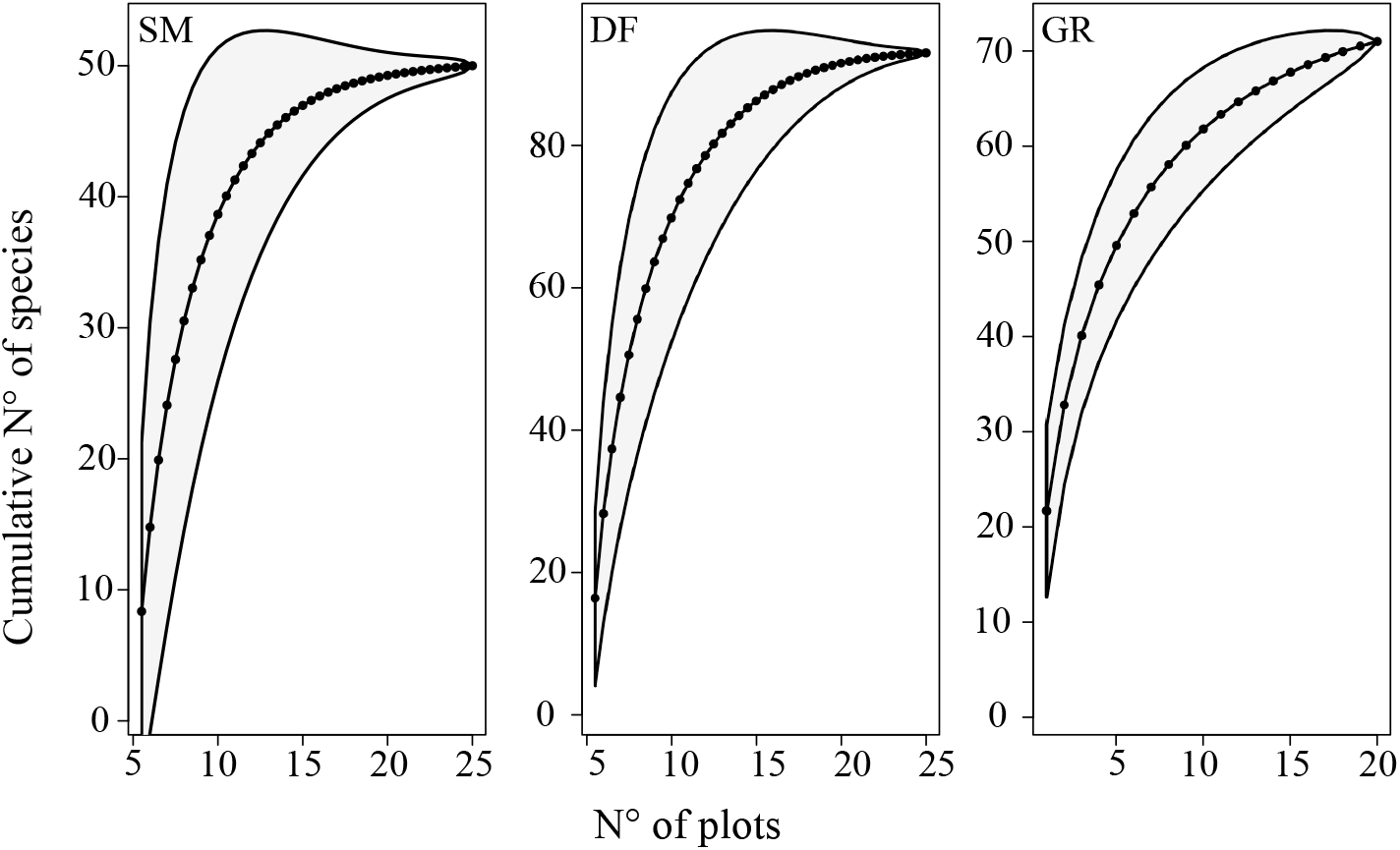
Species accumulation curves from the sampling of plants. Each curves represent a habitat (SM = Spruce Monoculture, DF = Deciduous Forest and GR = Grassland.

#### 6.2 B - Soil parameters

The supplementary soil parameters we measured are reported in following graphs (Figure 9).

**Fig. 9:**
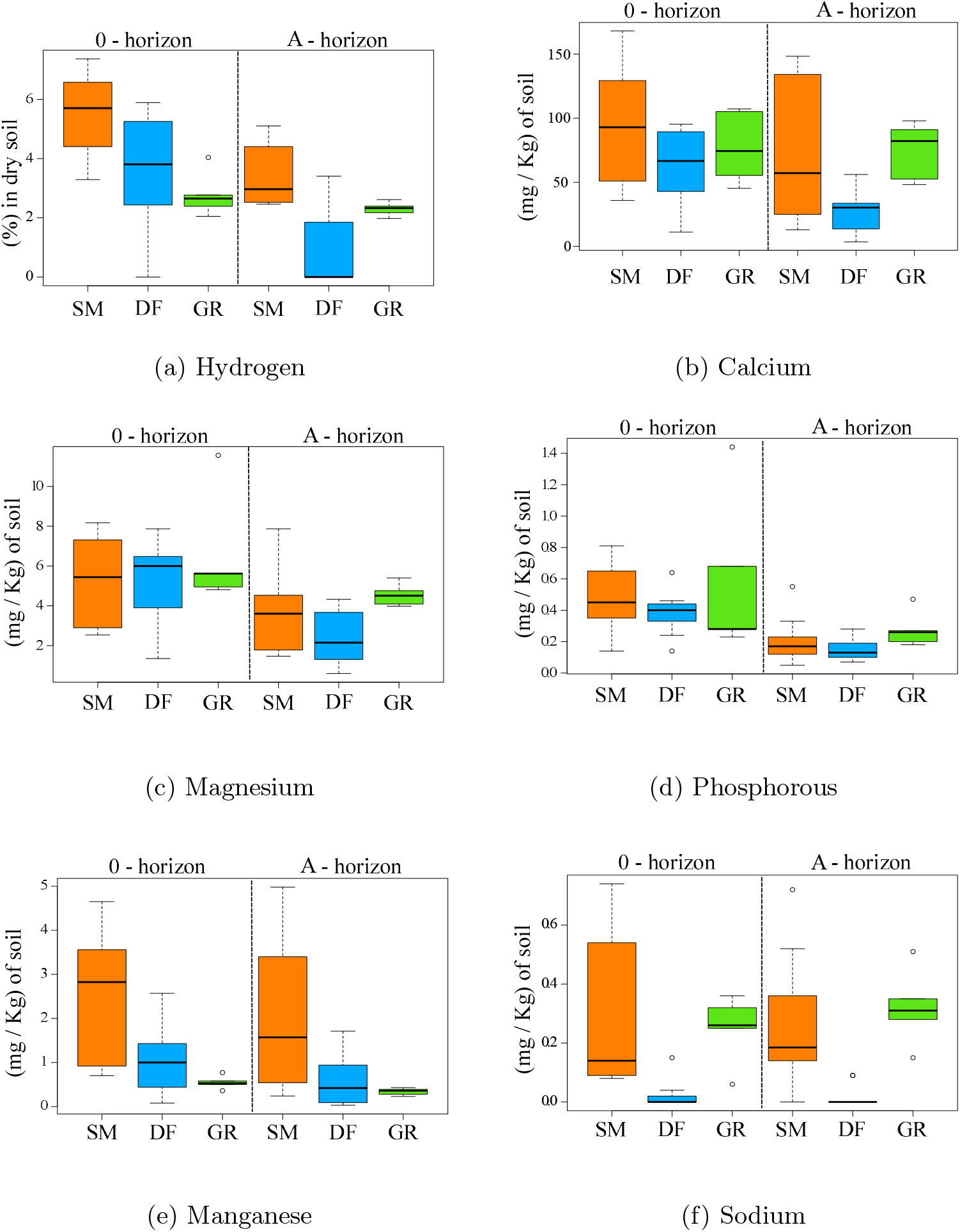
Box plots representing the soil parameters we did not considered in our analysis. On the x-axis are indicated the habitats (SM = spruce monoculture, DF = deciduous forest and GR = grassland). On the top of graphic are indicated the soil horizons (“0” horizon = litter, “A” horizon = organic mineral complexes). The y-axis shows the scales of each parameters taken into account.

